# Viruses encode tRNA and anti-retron to evade bacterial immunity

**DOI:** 10.1101/2023.03.15.532788

**Authors:** Aa Haeruman Azam, Kotaro Chihara, Kohei Kondo, Tomohiro Nakamura, Shinjiro Ojima, Azumi Tamura, Wakana Yamashita, Longzhu Cui, Yoshimasa Takahashi, Koichi Watashi, Kotaro Kiga

## Abstract

Retrons are bacterial genetic retroelements that encode reverse transcriptase capable of producing multicopy single-stranded DNA (msDNA) and function as antiphage defense systems. Phages employ several strategies to counter the host defense systems, but no mechanisms for evading retrons are known. Here, we show that tRNA^Tyr^ and Rad (retron anti-defense) of T5 phage family inhibit the defense activity of retron 78 and a broad range of retrons, respectively. The effector protein of retron 78, ptuAB, specifically degraded tRNA^Tyr^ leading abortive infection, but phage countervailed this defense by supplying tRNA^Tyr^. Rad inhibited retron function by degrading noncoding RNA, the precursor of msDNA. In summary, we demonstrated that viruses encode at least two independent strategies for overcoming bacterial defense systems: anti-defense, such as Rad, and defense canceler, like tRNA.

---

The retron defense system composed of reverse transcriptase (RT), non-coding RNA, msrmsd, and accessory protein or RT-fused domain with various enzymatic functions^1–3^. The RT produces satellite msDNA molecules using msd RNA as the template^4^. Following the production of msDNA, the msd RNA template is digested by RNase H^5^. The final product is typically a branched DNA-RNA hybrid in which msd DNA and msr RNA are covalently joined via a 2’-5’ phosphodiester bond^4^. In some cases, such as retron Ec78, Ec83 and Sen2, the msd DNA is further separated from the msr RNA^6,7^ by the housekeeping exonuclease VII encoding genes *xseA* and *xseB* ^6,7^. There are 13 different types of retrons based on their genetic structure and accessory proteins^8^. The accessory protein, which is hugely diverse across different retrons^8^, is the executor (effector protein) in retron defense that acts to abort phage infection through the inactivation of bacterial growth. In response to anti-phage defenses, phages have developed various counteract strategies, one of which is to encode proteins that inactivate defense, including these very recently identified anti-BREX^9,10^, anti-CBASS^11,12^, anti-Pycasr^11^, and anti-Thoeris^13^. The arms race between bacteria and phages is the natural driving force of the incessant emergence of sophisticated anti-phage defense systems whose discoveries and mechanistic understandings have brought about multiple impactful modern biotechnological tools.

## Phage genes that inhibit retron function

We have previously isolated and characterized a broad host range *Escherichia coli* phage ΦSP15^14^ that shares high similarity with T5j phage, a wildtype T5 from phage collection of Jichi Medical University. ΦSP15 was allowed to undergo spontaneous mutations through passage co-culture with bacteria under Fosfomycin addition. We identified one resultant mutant from each phage - T5n and ΦSP15m, respectively, each carries an approximately 8kb-deletion region in their genome, which is later found to encode multiple anti-defense systems, and we denoted <u>A</u>nti-<u>D</u>efense <u>I</u>sland (ADI) (Fig 1A). The mutant T5n is a T5 strain obtained from Biological Research center, National Institute of Technology and Evolution (Tokyo, Japan) that may have undergone mutation during routine propagation. We evaluated the ability of these four phages to infect a bacterial library encompassing different types of antiphage defense system^3^. Both deletion mutants, T5n and ΦSP15m, showed significant reduction in their infectivity against bacteria carrying Retron Ec67 and Retron Ec78 (Fig 1B and 1C, extended Fig 1). The ADI from ΦSP15 was then divided into nine fragments, each separately cloned into plasmid pKLC23^15^ carrying pBAD inducible promoter and transformed into *E. coli* DH10B cells expressing Retron Ec67 or Retron Ec78, revealing that fragment 8 (F8 ADI) could rescue T5n and ΦSP15m from both Retrons whereas fragment 6 (F6 ADI) only rescued phages from Retron Ec78 (Fig 1, D-F, extended Fig 3, A-B, extended Fig 4, A-B). Additionally, we found that F7 ADI and F8 ADI provided protection to phages from another retron that was not used during the first screening, Retron Ec83 (Fig 1F and 1I, extended Fig 5E). We discovered that ORF75 of ΦSP15 which we named rad (retron anti-defense) was the genetic determinant in F8 ADI that enabled the phage to evade the three retrons. Meanwhile, the tRNA^Tyr^ in F6 ADI was responsible for phage rescue from Ec78, and ORF71 and ORF72 of F7 ADI rescued phage from Ec83 (Fig 1, G-I. extended Fig 3c, Extended Fig 4, C-D and F-G, Extended Fig 5).

**Figure 1.**
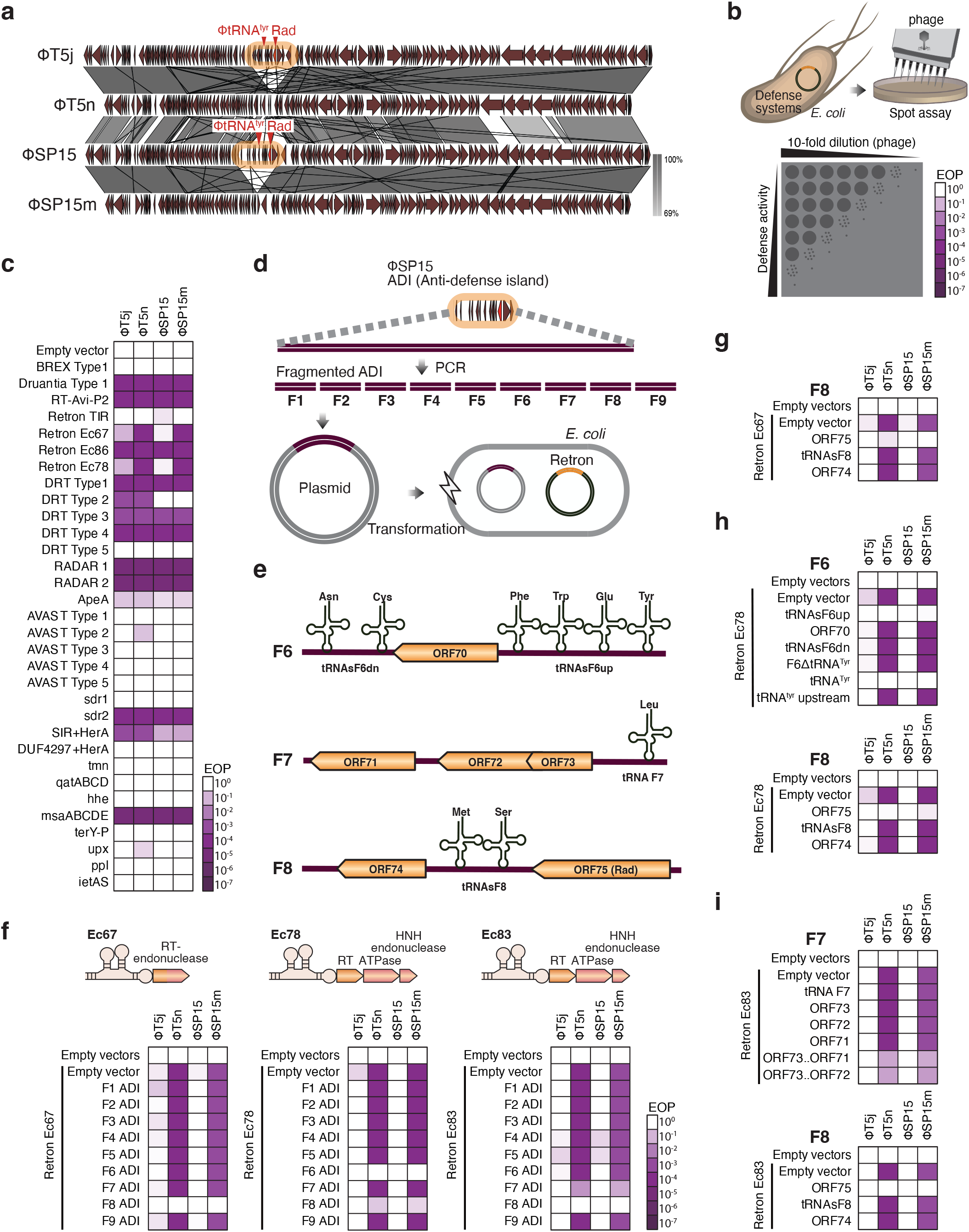
Identification of phage genes involved in retron evasion. (a) Genomic comparison of T5 and T5-like phage SP15. A genomic region of approximately 8 kb hereafter denotes as <u>A</u>nti-<u>D</u>efense <u>I</u>sland_(ADI) in T5j and SP15 were missing in T5n and SP15m, respectively. The visualized genomic comparison was generated using Easyfig^1^. (b) Simplified depiction of phage spot assay to evaluate the phage infectivity against bacteria with different antiphage defense systems. Phage solutions from serial 10-fold dilutions were dropped on bacteria lawn and the efficiency of platting (EOP) was measure accordingly. (c) Heatmap depicting the EOP change based on spot assay of phages on bacteria carrying plasmid with different defense systems. The bacterial strain used in this assay was *Escherichia coli* DH10B, the plasmids with antiphage defense systems are provided by Feng Zhang^2^ and are available on Addgene. T5n and SP15m showed decreased EOP comparing to their respective wild-type T5j and SP15 on bacteria with retron Ec67 and Ec78 defense systems. T5n has decreased EOP comparing to T5j on bacteria carrying AVAST 2. (d) Fragmentation of ADI into nine fragments. The ADI fragments were separately cloned into plasmid under a pBAD inducible promoter and co-transformed with retron into *E coli* DH10B. (e) Genetic organization of fragment 6 (F6 ADI), fragment 7 (F7 ADI), and fragment 8 (F8 ADI). (f) Heatmap based on spot assay of phages on bacteria carrying retron and different ADI fragments. F8 ADI neutralized defense activity of three different retrons Ec67, Ec78, and Ec83, tested in this panel. F6 and F7 ADI specifically neutralized Ec78 and Ec83, respectively. (g, h, i) Heatmap based on spot assay of phages on bacteria carrying retron and different F6, F7, and F8 ADI fragments. ORF75 in the F8 ADI neutralized all retrons tested, hereafter we name it <u>R</u>etron-<u>a</u>nti <u>d</u>efense (Rad). (h) In F6 ADI, tRNA^Tyr^ was found to be the genetic determinant responsible for Ec78 neutralization. The neutralization of F6 ADI was defective when tRNA^Tyr^ was deleted, and co-expression of tRNA^Tyr^ alone with Ec78 neutralized retron Ec78 defense activity. (i) Fragmentation of F7 ADI. Co-expression of two ORF72 and ORF73 are necessary to neutralize retron Ec83. Neither tRNA nor any single gene from F7 could neutralize Ec83. Empty vector pLG001 or pSC101 was used as negative control in all spot assay performed in this figure. Empty vector**<u>s</u>** indicate co-expression of empty vector and plasmid carrying retron.

## Rad degrades retron ncRNA

Rad is a small protein (189 amino acids) of unknown function, with primase/helicase and TOPIRM/RNase domain (Supplementary Table S1). A search based on homology identified 541 Rad homologues in 19,263 phage genomes in the NCBI database. Rad-encoding phages belong to two families; *Siphoviridae* and *Myoviridae* and infect at least nine different genera of bacteria from three taxonomic phyla; *Proteobacteria, Cyanobacteria*, and *Actinobacteria* (Fig 2A, supplementary Table S2). We demonstrated that the Rad homolog from *Proteus mirabilis* phage (Rad^Proteus phage Privateer^) could strongly protect phages from Retron Ec78, while other Rad from *Shigella sonnei* phage (Rad^Shigella sonnei phage^) and Salmonella phage vB_Sen_I1 (Rad^Salmonella phage vB_Sen_I1^) showed moderate protection (Fig 2B and C, extended Fig 6A). Notably, Rad exhibited extensive inhibition against retrons including retron Ec48 and Se72 (Fig. 2D, Extended Fig 7)^1^. The anti-retron activity of Rad showed significant decrease by the introduction of single amino acid mutations at various locations that were conserved in other Rads (R13E, P33T, I88T, D135H, and E156H) (Fig 2, E-F, Extended fig 6B, Extended Fig 8, A-B). And when double mutations at selected conserved amino acids were introduced, Rad defense activity was completely abolished regardless of the location (Fig 2F, extended Fig 8C). Rad was shown to reduce msDNA and ncRNA (msr-msd transcriptional cassettes) of retron, but not the transcript of RT and effector protein, indicating that Rad may degrade the ncRNA to prevent further synthesis of retron (Fig 2, G-H). Exogenous expression of Rad via genome insertion improved T7 infectivity to bacteria carrying retron Ec67 (Fig 2I, Extended Fig 9).

**Figure 2.**
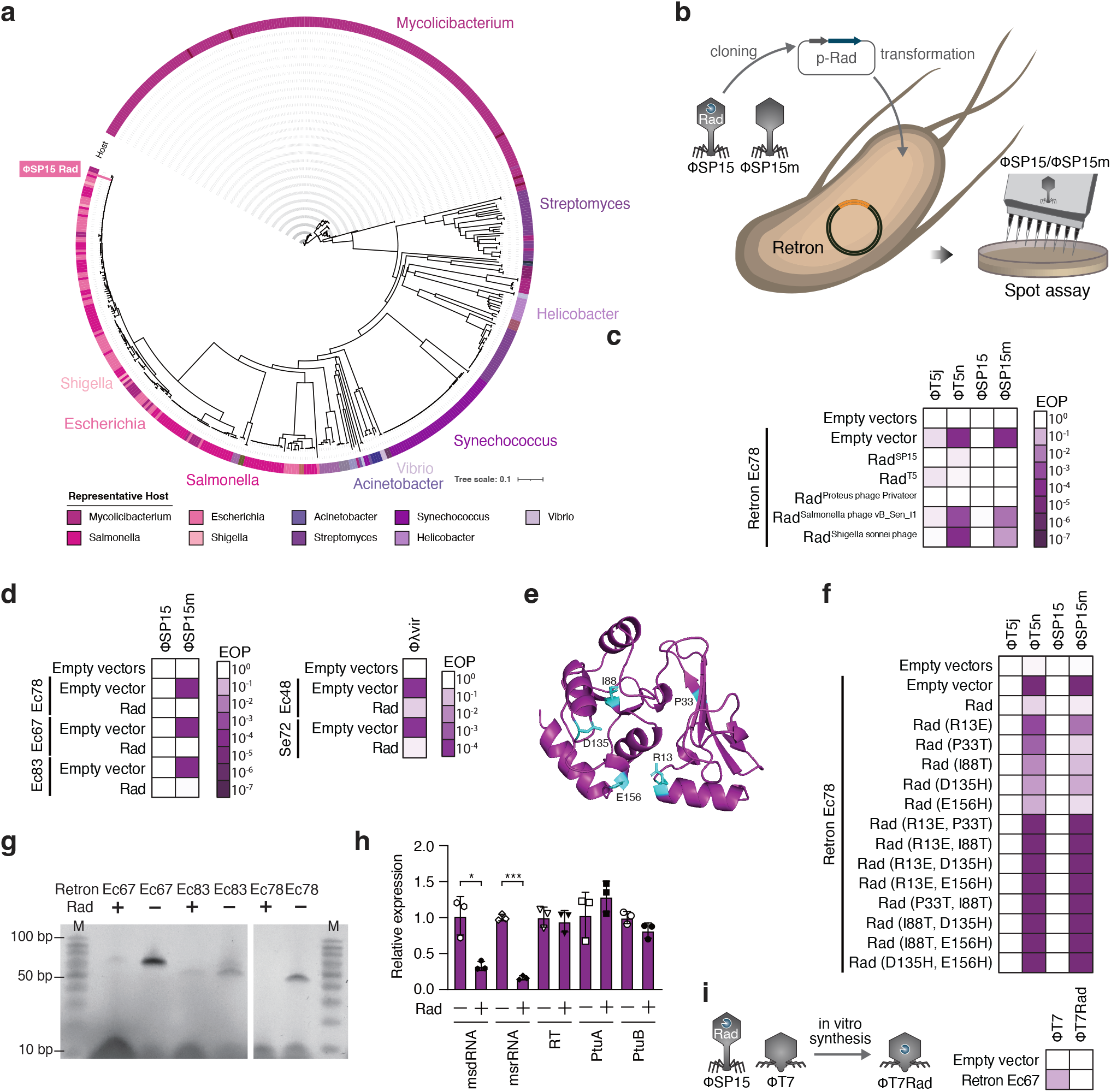
Rad is a potent blocker of retron defense that is widespread in phage infecting distinct genera. (a) Phylogenetic tree of Rad homologues from diverse bacteria genera. Bacteriophages that infect at least nine genera of bacteria carry Rad homolog, among these the Mycolicibacterium phages made up 40% of the Rad-carrying phages. (b) Simplified depiction of co-expression of Rad and retron. Rad was cloned into plasmid with pBAD inducible promoter. Spot assays were conducted using wildtype phage SP15 or T5j and their corresponding 8 kb deletion mutants, T5n or SP15m. (c) Rad from different phages that infected distinct taxa was capable of neutralizing retron. Rad from Salmonella phage vB_Sen_I1 and *Shigella sonnei* phage only slightly impair retron defense, whereas Rad from SP15, T5, and proteus phage Privateer all demonstrated substantial inhibitory action. (d) Rad blocks different retrons of distinct group. Rad effectively blocked retron Ec67, Ec78, and Ec83 and rescued SP15m. Utilizing phage *λ* vir, Rad rescued the phage from retron Ec48 and Se72. (e) Predicted structure of Rad using Alphafold. Amino acid mutations were introduced into conserved residues in Rad at different locations. (f) Single amino acid mutation in all selected locations slightly reduced activity of Rad to block retron. Rad activity was completely hampered when each of the selected amino acid mutant was introduced together. (g) TBE-Urea PAGE of extracted msDNA from bacteria expressing retron. The msDNA product was significantly reduced when Rad was co-expressed with retron. Same result was observed in three different retrons Ec67, Ec78, and Ec83. (h) Real time quantitative PCR of retron cassette (msr-msd, RT, and effector protein) of Ec78. The relative expression of msr and msd RNA were significantly lower when Rad is co-expressed with retron. The experiment was conducted on three independent samples. Asterix indicates significant difference (\*\*\**P*<0.01, \**P*<0.05, according to student t-test). (i) Exogenous expression of Rad in T7 phage. Chimera T7 carrying *rad* gene (T7rad) exhibited increased infectivity by tenfold against bacteria carrying retron Ec67.

## ptuAB of retron Ec78 degrades tRNA^Tyr^

Since the inhibition of Retron by tRNA^Tyr^ was specific to Ec78 (Fig. 1F), we firstly focused on the effector proteins, which show the highest variations in the retron gene cluster. Retron Ec78 has two effector proteins, PtuA with an ATPase domain and an HNH endonuclease PtuB^1,3^. We then expressed the effector proteins individually (PtuA or PtuB) or together (PtuAB) under the inducible promoter pBAD. We demonstrated that PtuAB, but not the singly expressed effectors, triggered bacterial growth arrest, indicating that PtuA and PtuB are the toxins of retron Ec78 (Fig 3, A-B). When RT was removed from Retron Ec78, PtuAB toxicity was observed, but not when msrmsd was eliminated, indicating that the antitoxin activity against PtuAB requires RT alone (Fig 3B). This mode of action is different from the tripartite toxin-antitoxin observed in Retron Sen2. RNA hybridization assay showed that the bacterial tRNA^Tyr^ was significantly depleted by PtuAB overexpression (Fig 3, C-D, extended Fig 10A). tRNA sequencing then confirmed that both bacterial tRNA^Tyr^, tRNA^TyrU^ (tRNA-Tyr-GTA-2-2) and tRNA^TyrV^ (tRNA-Tyr-GTA-1-1), were specifically down regulated in the bacteria where PtuAB expression was induced (Fig 3E, extended Fig 10, B-D). Taken together, these results reveal Retron Ec78 exerts its defense mechanism by aborting phage infection through depletion of bacterial tRNA^Tyr^ via PtuAB.

**Figure 3.**
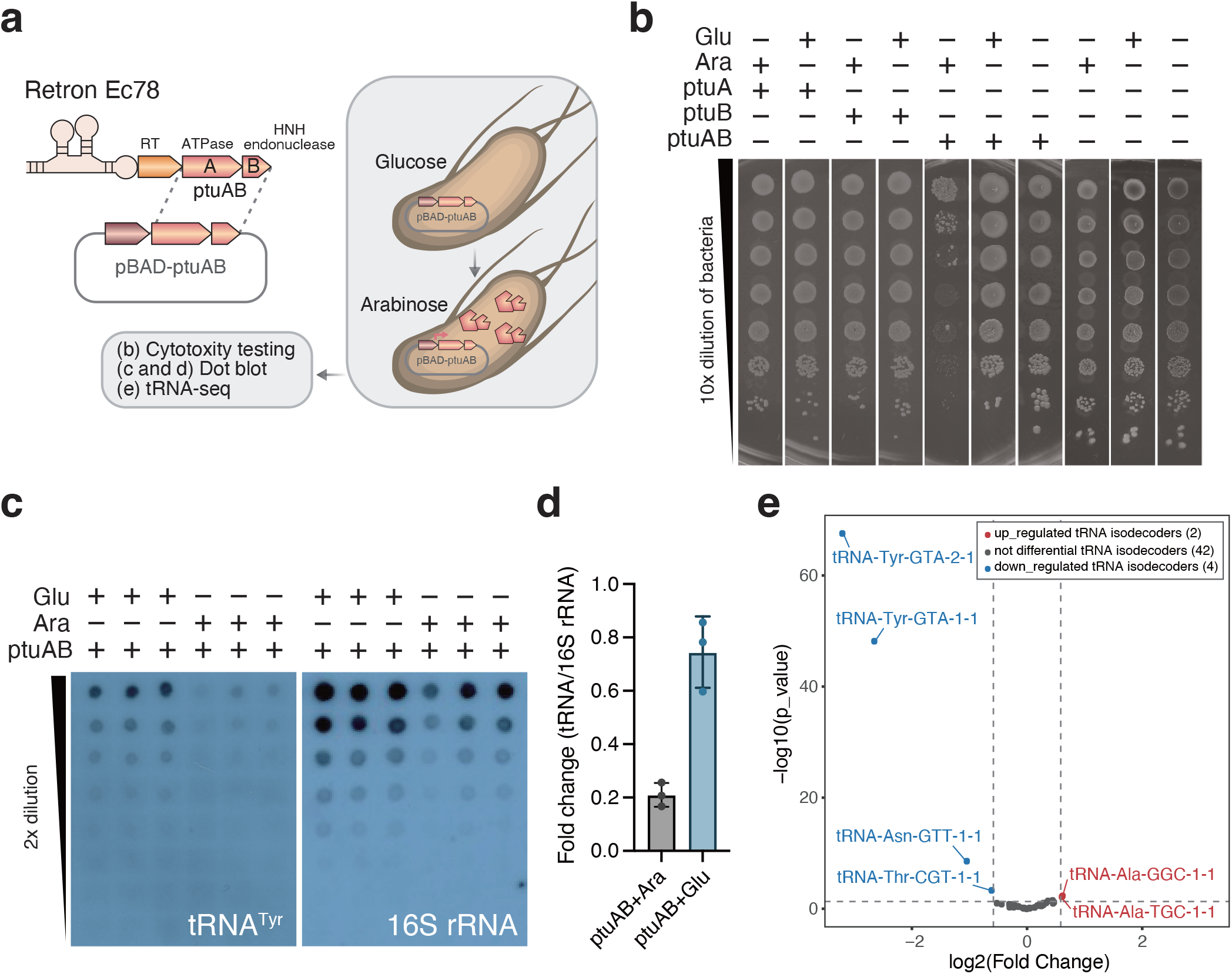
tRNA^Tyr^ is the cellular target of retron Ec78’s effector proteins. (a) Simplified depiction of method to evaluate cellular target of PtuAB of Ec78. The effector protein of Ec78 (PtuAB) was cloned individually, PtuA or PtuB, or together, PtuAB, into plasmid under pBAD promoter. Glucose was used to block pBAD promoter while arabinose was used to induce the promoter. The induced cell was evaluated for their cytotoxicity (b), and the reduction in the expression of tRNA^Tyr^ by dot blot RNA hybridization (c and d) and tRNA sequencing (e). (b) Induction of PtuAB promotes bacterial growth arrest. Induction of PtuA or PtuB alone was not toxic to bacteria, whereas induction of both (PtuAB) was toxic. (c and d) tRNA^Tyr^ were significantly low in the bacteria where PtuAB was expressed. The intensity of the dot obtained from the RNA hybridization assay was visualized using ImageJ (d). (e) tRNA sequencing revealed the tRNA^Tyr^ was significantly down regulated when PtuAB was induced.

## Phage tRNA^Tyr^ cancels abortive infection

Because phage derived tRNA^Tyr^ (ΦtRNA-Tyr_SP15) in F6 ADI can rescue phages from Retron Ec78 (Fig 1H), we speculated that ΦtRNA-Tyr_SP15 neutralizes Retron Ec78 through a different mechanism than Rad. Since changing the anticodon sequence of ΦtRNA-Tyr_SP15 or mutating the stem-loop sequence of ΦtRNA-Tyr_SP15 exterminated the neutralization effect of ΦtRNA-Tyr_SP15 (Fig 4, A-E, Extended Fig 11A), we presumed that the function of ΦtRNA-Tyr_SP15 in protein synthesis would be essential for the inhibition of retron defense.

**Figure 4.**
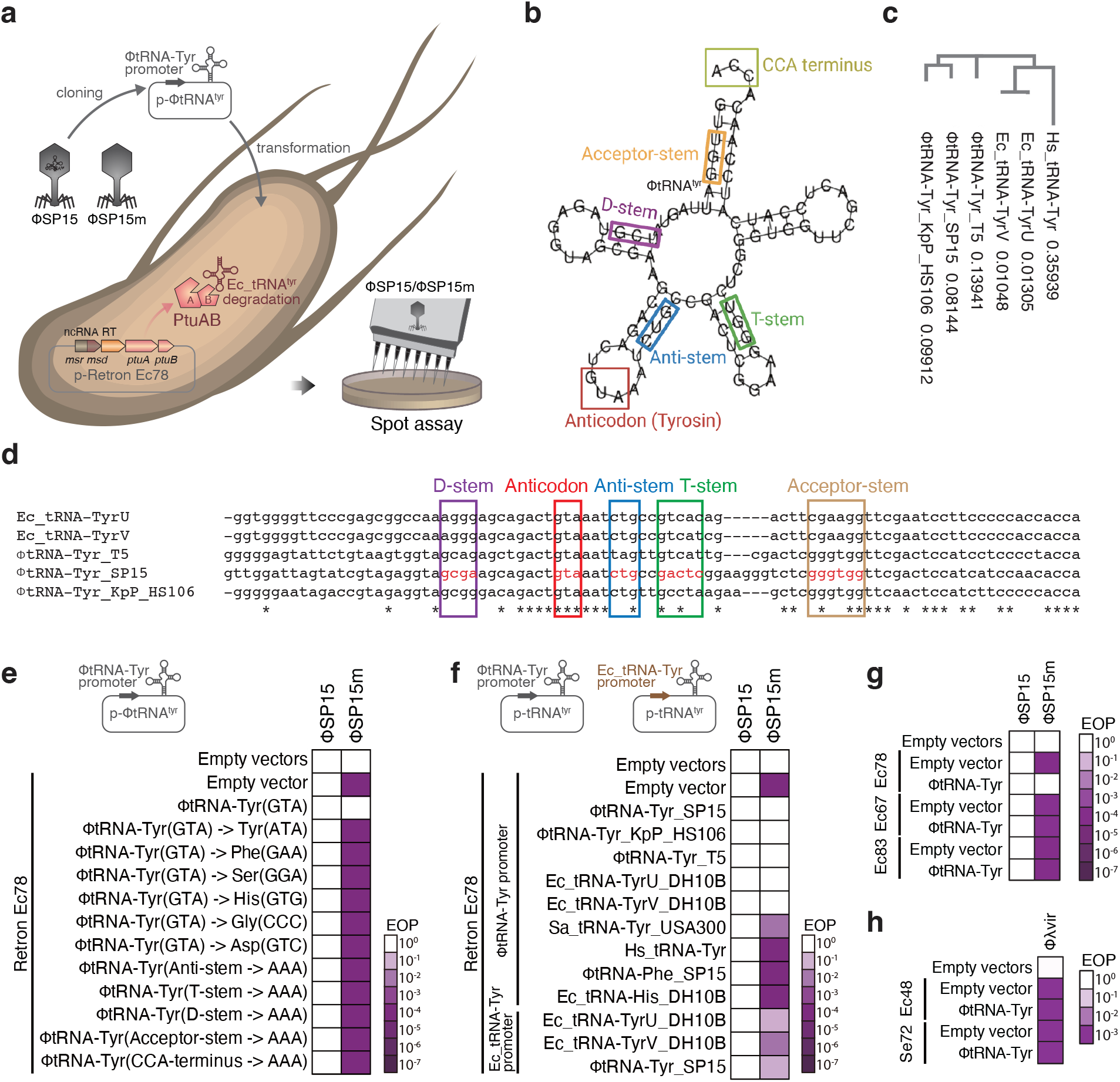
tRNA^Tyr^ from other phages or from host bacteria could rescue phage from retron Ec78. (a) Simplified depiction of the method to evaluate tRNA complementation on bacteria carrying retron Ec78. Complementation of tRNA was performed *in trans* by cloning the tRNA into plasmid under phage SP15 derived tRNA promoter (ΦtRNA-Tyr promoter) located in F6 ADI. (b) RNAFold^3^-based structural prediction of tRNA^Tyr^ SP15 (ΦtRNA-Tyr_SP15). To assess the impact of the mutation on the restoration of the phage from retron Ec78, sequence and structural mutations were introduced into tRNA. These mutations included the CCA terminus (CCA into AAA, yellow-green box), acceptor stem (UGG into AAA, orange box), D-stem (UGG into AAA, violet box), anti-stem (GUC into AAA, blue box), and T-stem (GGU into AAA, green box). (c) Phylogenetic tree of tRNA^Tyr^ used in this study. (d) Sequence alignment of tRNA^Tyr^ from T5 (ΦtRNA-Tyr_T5), SP15 (ΦtRNA-Tyr_SP15), *Klebsiella* phage KpP_HS106 (ΦtRNA-Tyr_KpP_HS106), and *E. coli* tRNATyr (Ec-tRNA_TyrU or Ec-tRNA_TyrV). According to the predicted secondary structure, the loop, stem, and anticodon sequence of SP15 were highlighted in red letters within the colored boxes. (e) Heatmap based on spot assay of phage SP15 and SP15m on bacteria carrying retron Ec78 complemented with different mutant of ΦtRNA-Tyr_SP15. Regardless of the mutation locations, tRNA^Tyr^ mutations eliminated the tRNA^Tyr^’s capacity to neutralize retron defense. (f) tRNA^Tyr^ from different phages (ΦtRNA-Tyr_T5 and ΦtRNA-Tyr_KpP_HS106) or from *E coli* rescue phage from retron Ec78. Notably, Ec_tRNA_Tyr could rescue the phage to the same extent as ΦtRNA-Tyr_SP15 when SP15 tRNA promoter was utilized, however when *E. coli* tRNA promoter was used (Ec_tRNA-Tyr promoter), the retron defensive activity was still visible. Other tRNAs (ΦtRNA-Phe__SP15 or Ec_tRNA-His_DH10B) and tRNATyr from human (Hs_tRNA_Tyr) or *Staphylococcus aureus* (Sa_tRNA_Tyr_USA3OO) did not rescue phage from retron Ec78. (g and h) tRNA^Tyr^ specifically rescued phages from retron Ec78. Co-expression of tRNA^Tyr^ with other retron was not able to rescue phage SP15m from retron Ec67, Ec78, nor Ec83 (g). Similarly, tRNA^Tyr^ was not able to rescue phage *λ* vir from Ec48 nor Se72 (h).

Complementation of the exogenous tRNA^Tyr^ by either T5 tRNA^Tyr^ (ΦtRNA-Tyr__T5), *Klebsiella* phage KpP_HS1O6 tRNA^Tyr^ (ΦtRNA-Tyr__KpP__HS1O6), and endogenous host bacteria *Escherichia coli* DH10B tRNA^Tyr^ (Ec_tRNA-TyrU or Ec_tRNA-TyrV) *in trans* under the SP15 derived tRNA promoter (ΦtRNA-Tyr promoter) successfully restored phage infection to that of ΦtRNA-Tyr_SP15, instead only partial recovery was observed by complementation under *E. coli* tRNA promoter (Ec_tRNA-Tyr promoter) (Fig 4F, Extended Fig 11B). The ability of tRNA^Tyr^ to rescue phage from retron defense was only observed on retron Ec78 (Fig 4, G-H, Extended Fig 11C).

## Phage sensing mechanism of Retron Ec78 and Retron Ec67

The retron defense system works by sensing phage infection and activating the effector protein(s). Various genetic determinants of phage that triggered antiphage defense systems have been studied elsewhere^16^, but only a small number of retron triggers have been identified. We sought to determined how the two retrons, Ec78 and Ec67, recognize and mitigate phage infection through screening for phage mutants that could bypass each of the retrons. T5n, ΦSP15m, and T2 were employed for Retron Ec67, while T5n and ΦSP15m were used against Retron Ec78 (Fig 5A).

**Figure 5.**
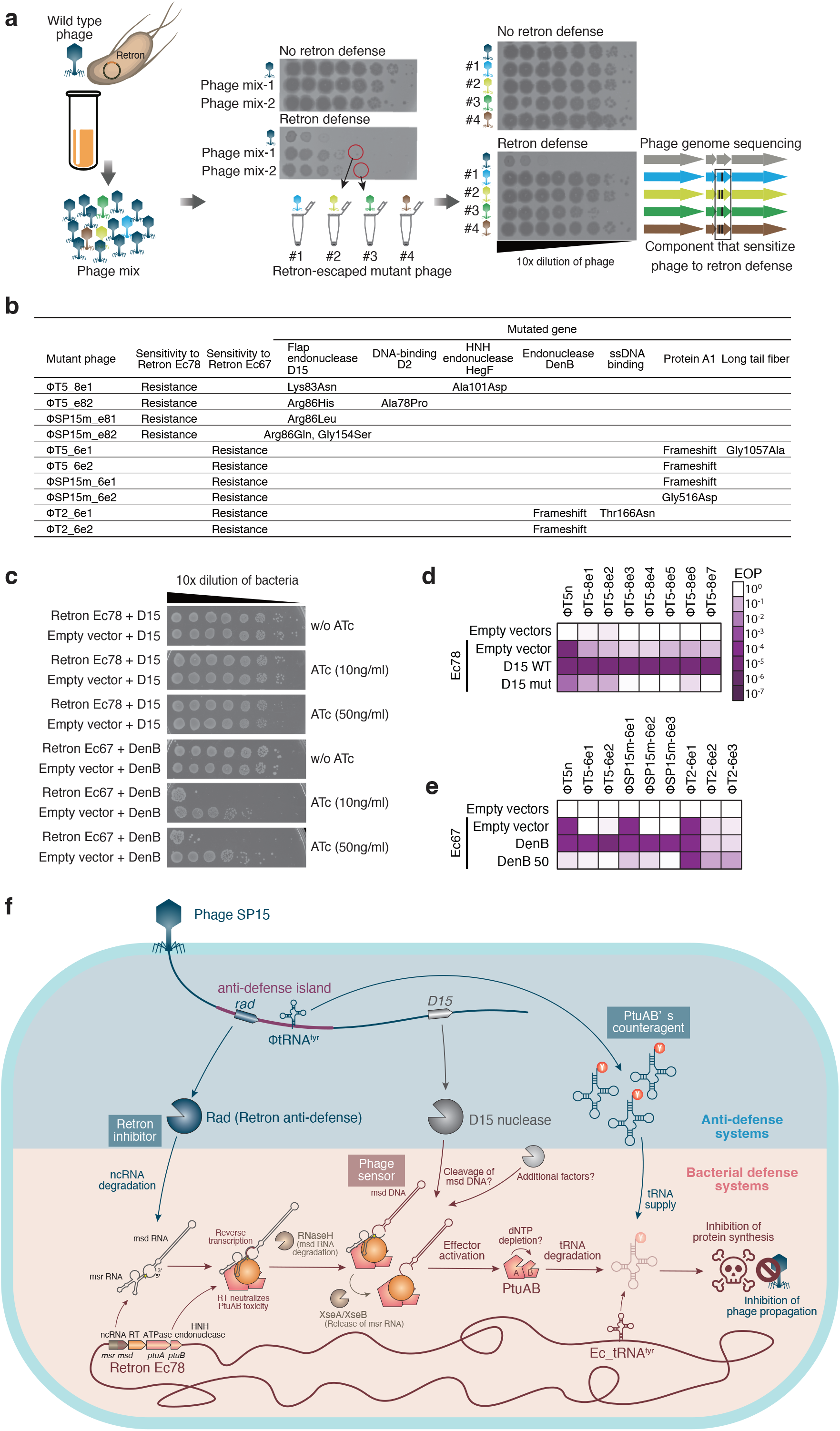
Evaluation of retron trigger. (a) Screening of retron escaping phages adapted from another study^4^ with some modifications. Phage and bacteria harboring retron were cocultured in liquid medium overnight. Escaping mutant phages were screened from phage mixture obtained from the co-culture. Such mutant phages should form single plaque even when retron is presented in bacteria. The genomes of selected escaping mutans were analyzed and mapped to parental phage genome. Shared mutations in all escaping phages are expected to be the phage component that desensitized the phage to retron defense. (b) Mutations identified in escaping mutant phages. (c) Phage genes that are commonly mutated in escaping phages were tested for their toxicity when co-expressed with retron. D15 protein that was mutated in all Ec78 escaping phages was not toxic even when co-expressed with retron Ec78. DenB protein from Ec67 escaping T2 phage was highly toxic when co-expressed with Ec67, suggesting the importance of the gene in the activation of Ec67. (d) Complementation of D15 restored retron defense against escaping phages, T5-8e andSP15m-8e, whereas complementation of the mutated version of D15 did not restore retron activity, indicating D15 may involve in retron Ec78 activation. (e) DenB complementation restored Ec67 defense against the escaping phages T5-6e, SP15m-6e, and T2-6e. Co-expression of DenB without ATc induction could restore retron defense against escaping phage T5-62 and SP15m-6e, but induction with 50ng/ml ATc (DenB 50) was needed to see the change in the escaping phage T2-6e. However, in such condition, retron Ec67 defense activity was not visible against escaping phage T5-6e and SP15m-6e, perhaps due the high toxicity of DenB. (f) Proposed mechanism of phage to escape retron Ec78. Phages encode three genetic factors related to retron defense; Retron-anti defense (Rad) that inhibited retron biosynthesis, retron trigger (D15 for Ec78, DenB or A1 for Ec67), and tRNAs to supplement the host tRNAs that was degraded by retron Ec78. Following infection of phage, retron may sense the phage protein that is either pre-made and packed together with phage genome in the capsid or the protein that is expressed during early production of phage particle. Production of such sensor/trigger may activate retron defense system either due to degradation of retron component (because most retron triggers identified in the current study are interestingly involved in nucleotide degradation) or retron senses degradation of host genomic DNA. Once the retron is activated, effector protein will be released from retron complex. We suggested that the depletion of dNTPs due to the overused of those for phage assembly may further trigger conformational change of PtuAB to its active form that cleaves bacterial tRNAs, resulting in bacterial growth arrest and aborts phage production. To evade retron defense, however, phage is equipped with Rad and/or, in case of Ec78, the phage encoded tRNAs as the counter-agents of PtuAB effector protein.

For Retron Ec78, we found several missense mutations in the gene encoding for exonuclease D15 in all escaper mutants of T5n (seven) and ΦSP15m (four) (Fig 5B). D15 protein catalyzes both the 5’-exonucleolytic and structure-specific endonucleolytic hydrolysis of branched-DNA molecules^22–24^. Co-expression of D15 protein and Ec78 was not toxic to bacteria, but it restored Ec78 defense activity against escaper mutants of T5n and ΦSP15m (Fig 5C, extended Fig 12, A-B). Our findings suggest that Retron Ec78 defense may be triggered by not just D15 protein but also other unknown factor.

For Retron Ec67, we found a single point mutation that distinguishes the mutant phages from their parental strains in all seven T5 mutants, four ΦSP15m mutants, and three T2mutants (Fig 5B). Since all escaper phages of T2 and T5n/ΦSP15m carry mutations in DenB and protein A1, respectively, we presumed that these genes are the genetic determinants that activate Ec67. Both protein A1 and DenB are involved in DNA degradation; with protein A1 responsible for the degradation of host DNA as well as the shutoff of host genes^17–19^, whereas DenB protein cleaves single-stranded DNA in a dC-specific manner, which may be lethal to host dC-containing DNA replication^20,21^. Co-expression of DenB and Ec67 significantly inhibited bacterial growth (Fig 5C). However, we could not observe the effect of protein A1 due to its toxicity^16^. Although T5n and ΦSP15m do not carry DenB homolog protein, DenB complementation restored Ec67 defense activity against not only T2 escaper mutants but also those of T5n and ΦSP15m (Fig 5C, extended Fig 12C). These findings imply that Rectron Ec67 defense may be triggered by the activity of *denB* gene rather than DenB protein itself.

## Discussion

The current study describes Retron Ec78’s defense mechanism and identifies the cellular target of its effector protein PtuAB. These two proteins are also found in Septu^25,26^, an antiphage defense system with unknown molecular mechanisms. ATPase-like domain has been found in another nuclease mediated anti-phage defense system Gabija^25,26^. The GbjA protein of Gabija system consists of ATPase-like domain and TOPRIM domain. The ATPase-like domain is strictly regulated by nucleotide concentration *in vitro*, meanwhile the GbjA is activated by the depletion of dNTPs during phage infections, which in turn activates the TOPRIM domain with its nuclease activity, causing bacterial death^27^. We hypothesized that PtuA employs a similar strategy, which may be activated by dNTP depletion during phage infection. This could also explain why D15 protein, which is mutated in Ec78 escaper mutant phages, could not activate retron in the absence of phage infection. The expression of either PtuA or PtuB alone is not toxic to bacteria, suggesting these two proteins are most likely working together to induce bacterial growth arrest.

Retron Ec83 also has the same PtuAB effector as Retron Ec78, but failed to protect the phage by tRNA^Tyr^ complementation, suggesting that it is likely targeting other tRNAs or nucleic acids. ADI region is moderately preserved in Tequintavirus (Extended Fig 1B). In SP15, it encodes Rad in the tRNA-rich region, and another retron blocker in F7 ADI that specifically inhibits Retron Ec83 (Fig 1F, extended Fig5). Moreover, the absence of ADI in T5n resulted in lower infectivity against the antiphage defense AVAST 2 (Fig. 1B, extended Fig 1A), indicating the existence of an anti-AVAST 2 in T5n’s ADI and supporting the notion that anti-defense genes generally co-localize in a genomic island. Despite the genetic similarity of T-even phages, retron Ec67 used in the current investigation exhibited considerable inhibition on the T2 phage but only moderate inhibition on other T-even phages such as T4 and T6. We hypothesized that T4 and T6 might have other retron blockers, which allow them to evade Ec67. The lack of a Rad homolog protein in those phages, however, suggests that the T-even phage’s retron blocker(s), if present, may be different from Rad.

First discovered in the 1950s, transfer tRNAs (tRNAs) have been found to play a vital role in the central dogma of molecular biology in all living systems^28,29^. Only one decade after its first discovery, bacteriophages are found to carry their own tRNAs^30^. tRNAs are found in bacteriophage genomes from various bacterial genera^31^, but their precise function has long been elusive. Several hypotheses have been proposed, the most well-known is codon compensation, in which codons rarely used by the host but required by the phage are supplemented by phage-encoded tRNAs. This hypothesis is supported by the observation that phage-derived tRNAs tend to correspond with codons that are highly used by phage-encoded genes^32,33^. Recent studies may have hinted at another function of phage-derived tRNAs where they were discovered to be used by phages to counteract the depletion of host tRNAs that occurs as a general response to phage infection^34,35^. Our data showed that neither phage T5 nor SP15 endured any detrimental effects from losing a significant number of tRNAs in the ADI region. In contrast, these phages are no longer able to infect bacteria that are protected by the Retron Ec78 defense, suggesting that the phage tRNAs are involved in evading bacterial defense. So far, there has been multiple reports about nucleases, such as VapC^36,37^, PrrC^38^, or RelE^39^, of the toxin-antitoxin system, which are also known to target tRNA and are activated by various stress responses, including phage infection^40^. Most bacteria encode multiple defense systems, with an average of ~5 systems per genome^41,42^. This could be one of the reasons why T5-like phages, which carry multiple types of tRNAs in their genome, exhibits exceptionally broad host range^14^.

## Acknowledgments

Funding: This work was supported by the Japan Agency for Medical Research and Development (grant No. JP21fk0108496 and JP21wm0325022 to KK, JP21gm1610002 to LC and KK), JSPS KAKENHI (Grant No. 21H02110 and 21K19666 to KK). The funders had no role in the study design, data collection and analysis, decision to publish, or preparation of the manuscript.

