## Extended figures for "Viruses encode tRNA and anti-retron to evade bacterial immunity"

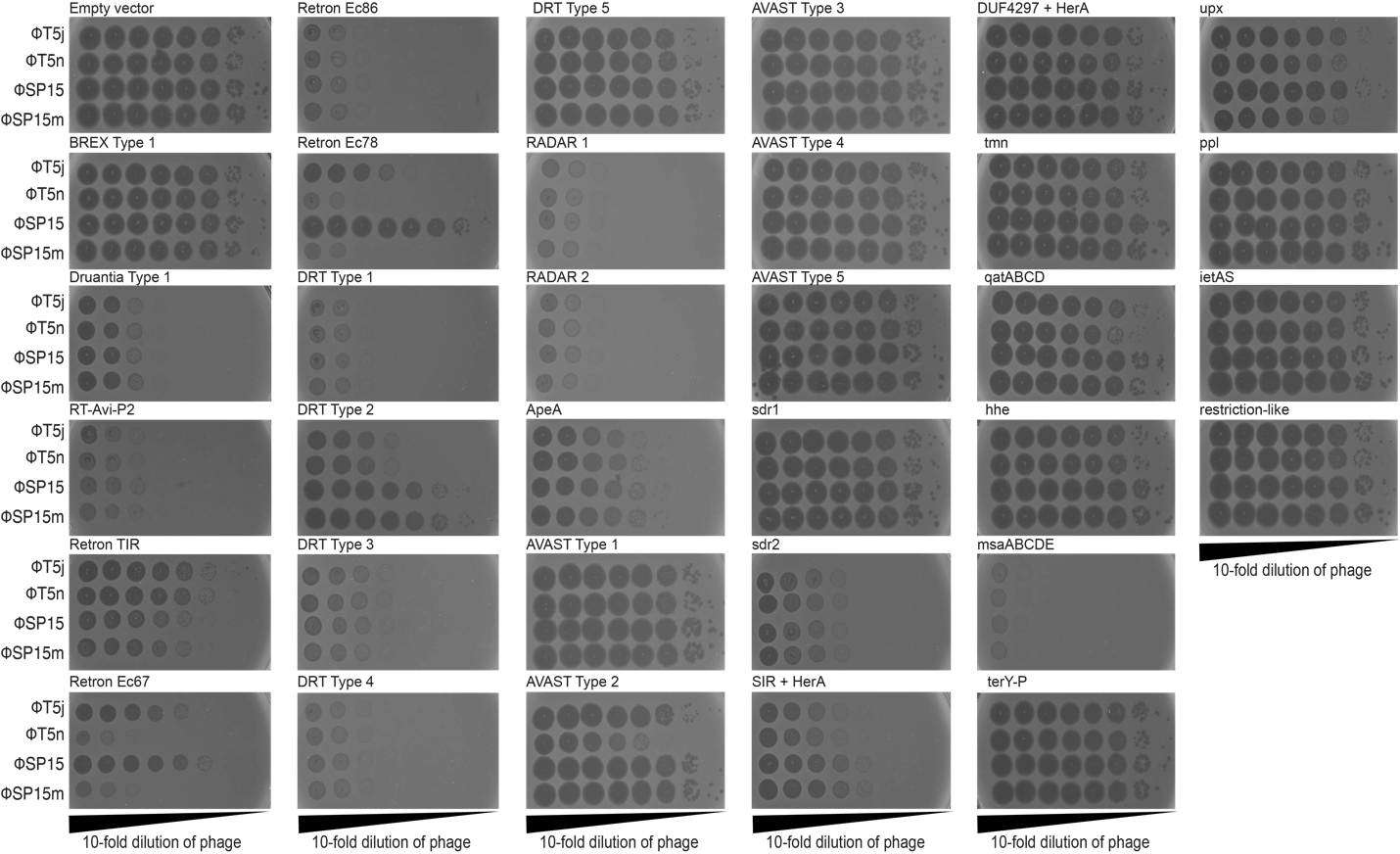

**Extended Figure 1.**

Spot assay of wildtype phages (T5j and SP15) and their respective mutants (T5n and SP15m) on bacteria carrying different antiphage defense systems from Gao et al^1^.

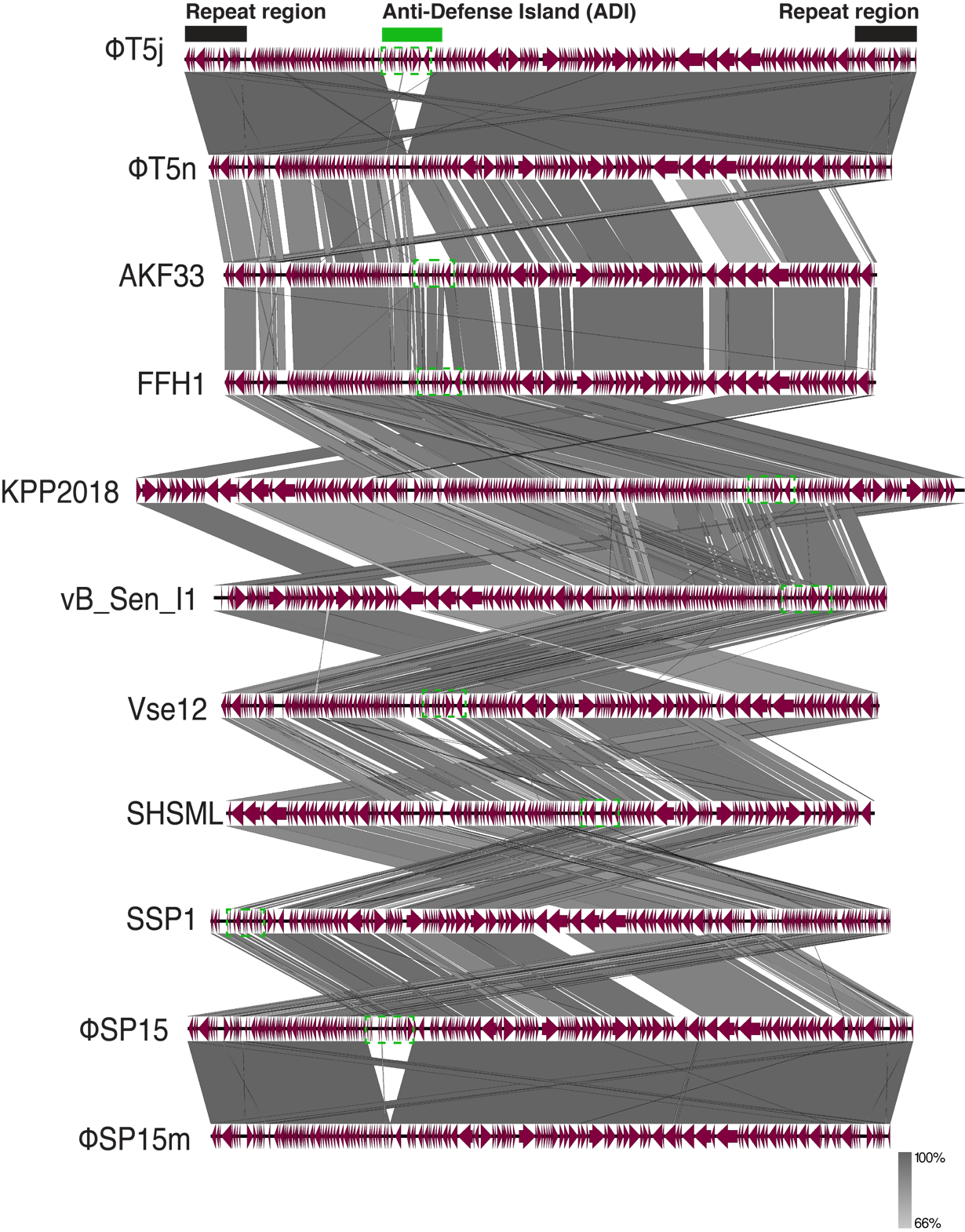

**Extended Figure 2.**

Conserveness of Anti-Defense Island (ADI) in T5-like phages infecting bacteria from various genera: T5-like *Escherichia coli* phages (T5j, T5n, SP15, SP15m, AKF33, and FFH1), *Klebsiella* phage (KPP2018), *Salmonella* phages (vB_Sen_I1 and Vse12), and *Shigella* phages (SHSML and SSP1). ADI regions are indicated by green dashline boxes.

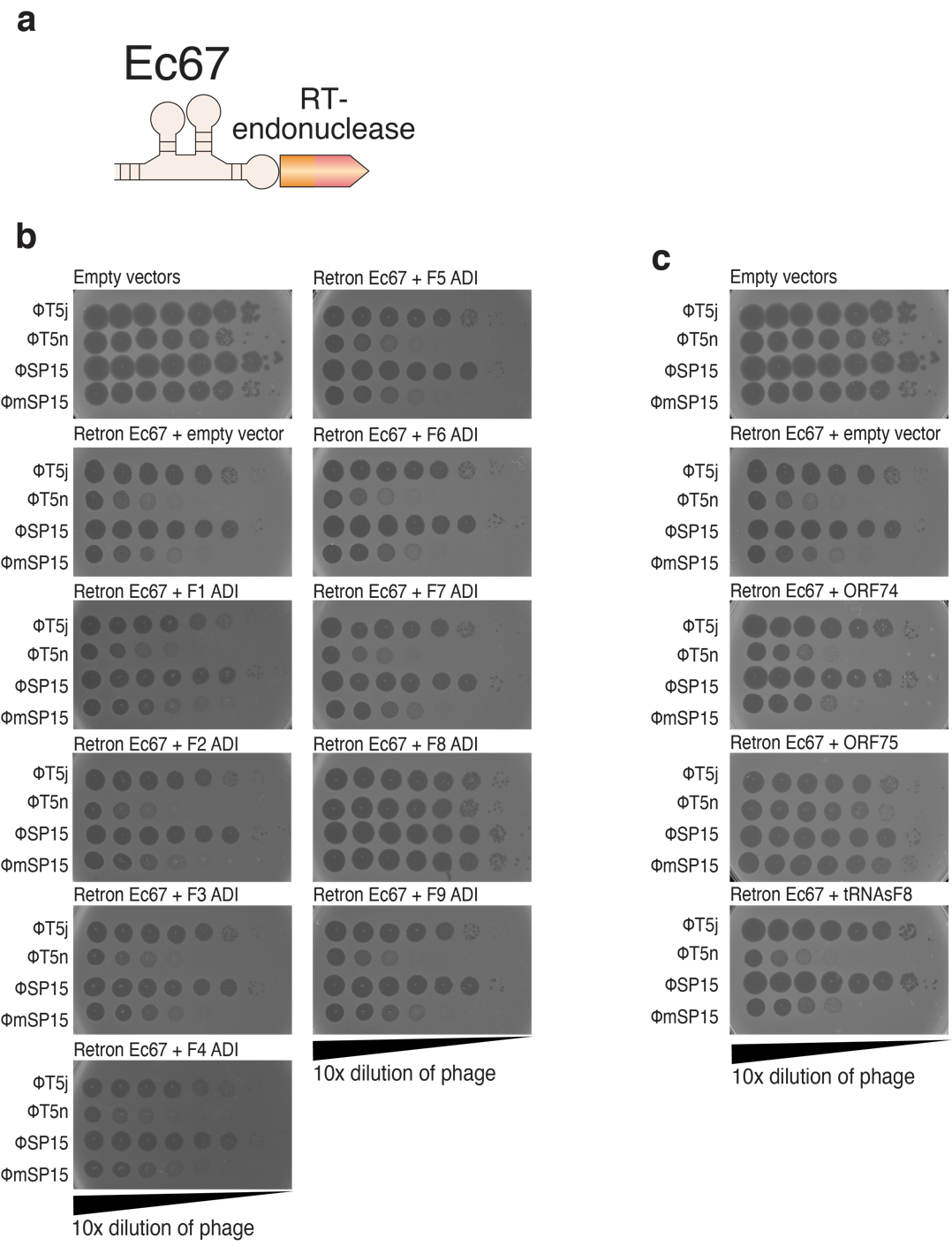

**Extended Figure 3.**

(a) Genetic organization of retron Ec67 consisting of non-coding RNA, and RT-fused effector protein with TOPRIM endonuclease domain. (b) Spot assay of wildtype phages (T5j and SP15) and their respective mutants (T5n and SP15m) on bacteria carrying retron Ec67 and different ADI fragments or different regions of F8 ADI (c).

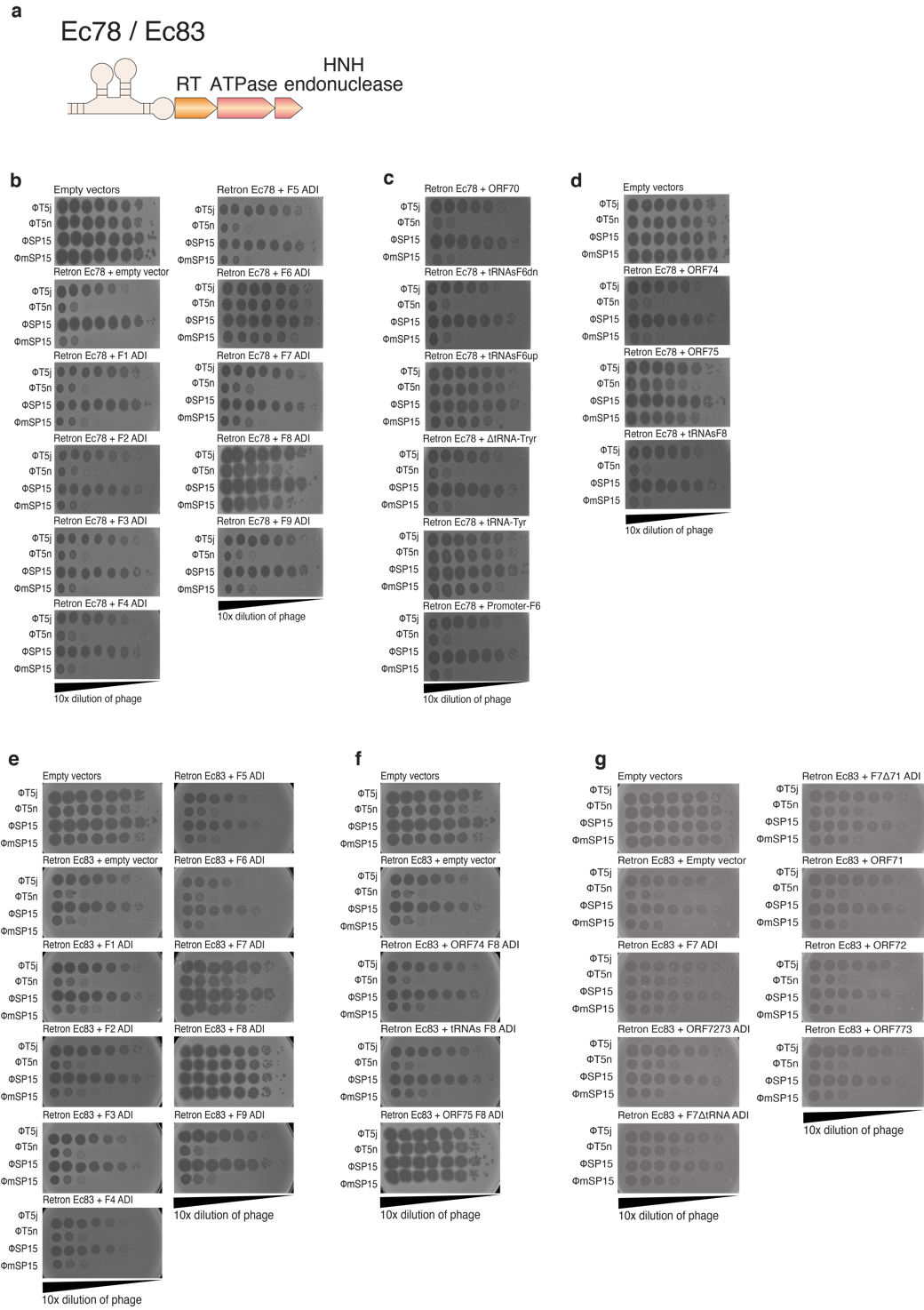

**Extended Figure 4.**

(a) Genetic organization of retron Ec78 and Ec83 showing non-coding RNA, reverse transcriptase (RT), and two effector proteins with ATPase (PtuA) and HNH Endonuclease domain (PtuB). Spot assay of wildtype phages (T5j and SP15) and their respective mutants (T5n and SP15m) on bacteria co-expressing retron Ec78 and different ADI fragments (b), or different regions of F6 ADI (c) and F8 ADI (d). Spot assay of phages (wildtype and mutants) on bacteria co-expressing retron Ec83 and different ADI fragments (e), or different regions of F7 ADI (f) and F8 ADI (g).

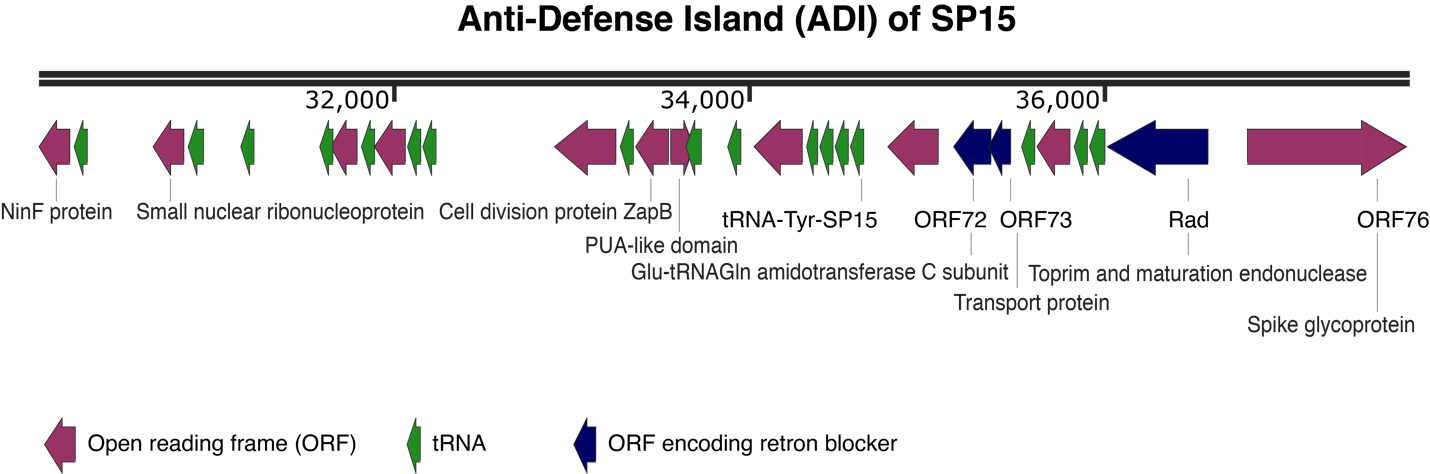

**Extended Figure 5.**

Genome map of the Anti-Defense Island (ADI) of SP15. Open reading frames (ORFs) are indicated by purple arrows, with the ORFs for retron blockers shown in blue. Green arrows indicate tRNAs. The numbers under the double black line designate genomic locations. The annotated proteins for each ORFs shown for each ORFs are matching hits with >= 50% probability from Pfam and COGS databases.

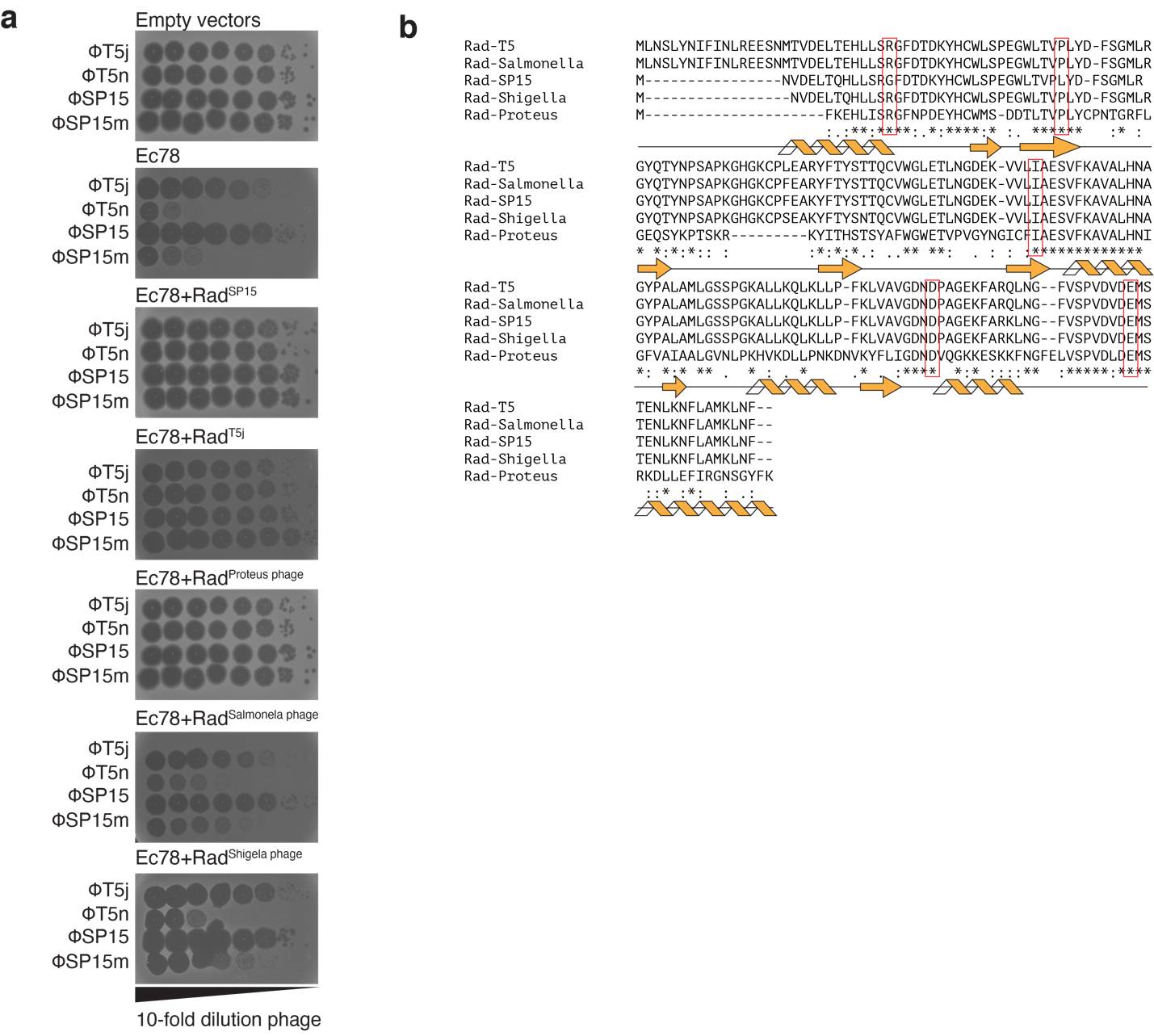

**Extended Figure 6.**

(a) Spot assay of wildtype phages (T5j and SP15m) and their respective mutants (T5n and SP15m) on bacteria co-expressing Rads from different phages and retron Ec78. (b) Protein alignment of Rads used in this study. The secondary structure of aligned proteins was shown based on structural prediction of Rad-SP15 using Alphafold. The amino acid residues that were selected for amino acid substitutions in Fig 2E and 2F are indicated by red boxes.

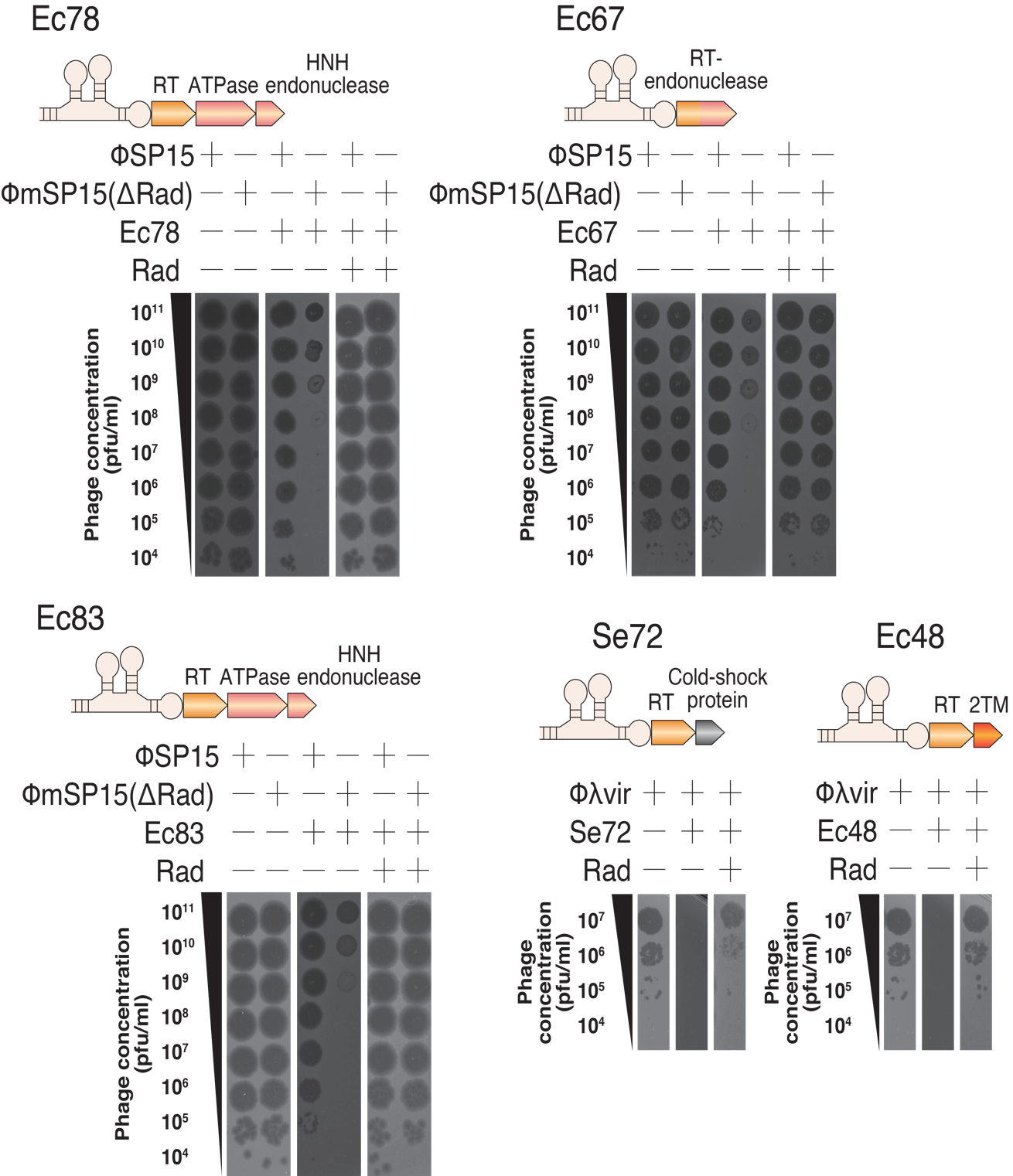

**Extended Figure 7.**

Spot assays of phages on bacteria carrying Rad-SP15 and different type of retrons.

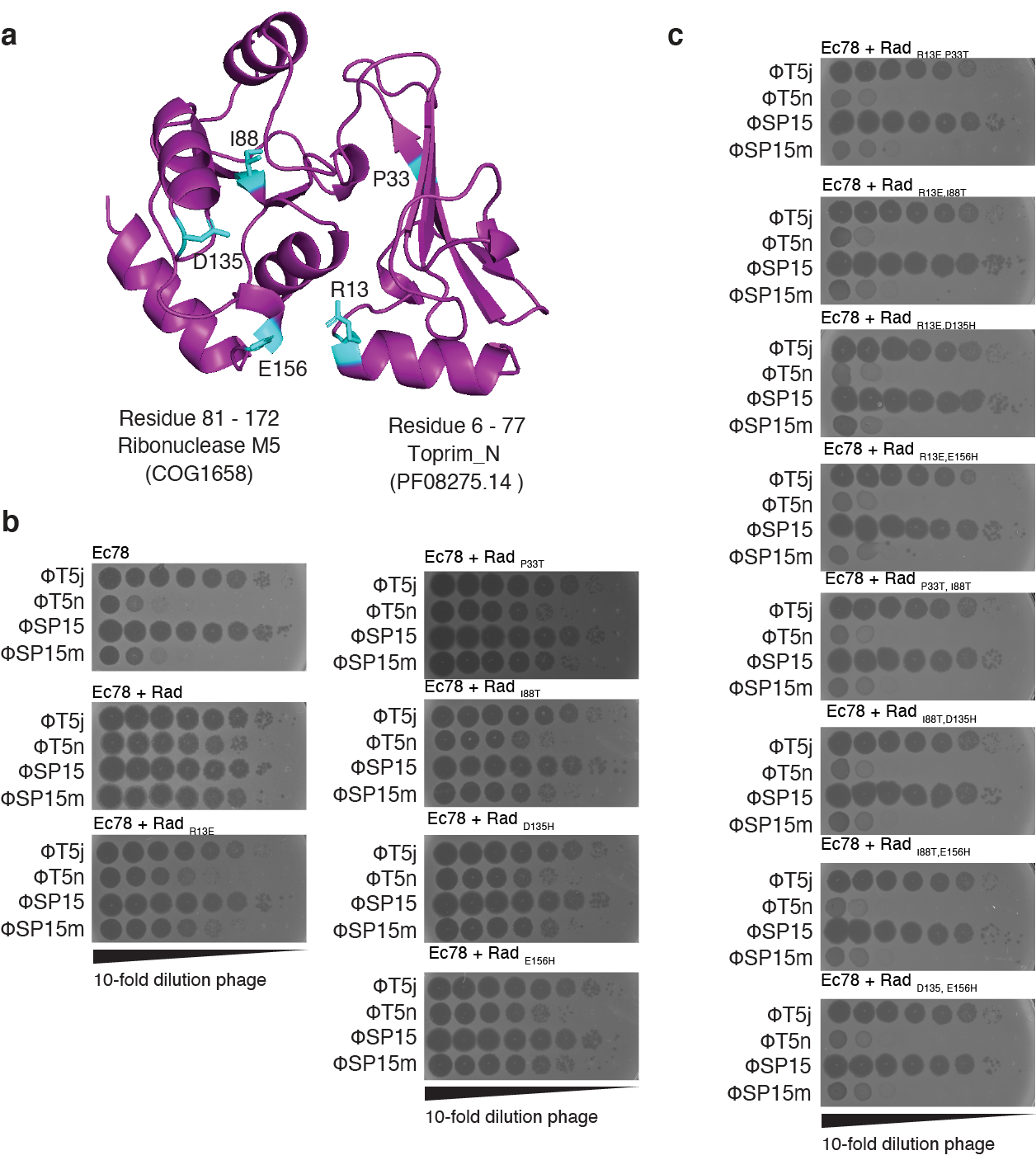

**Extended Figure 8.**

(a) Predicted structure of Rad-SP15 by Alphafold with the amino acids selected for mutation experiment and their positions highlighted. Spot assay of phages on bacteria that carry Rad with single amino acid mutations (b) and double amino acid mutations (c).

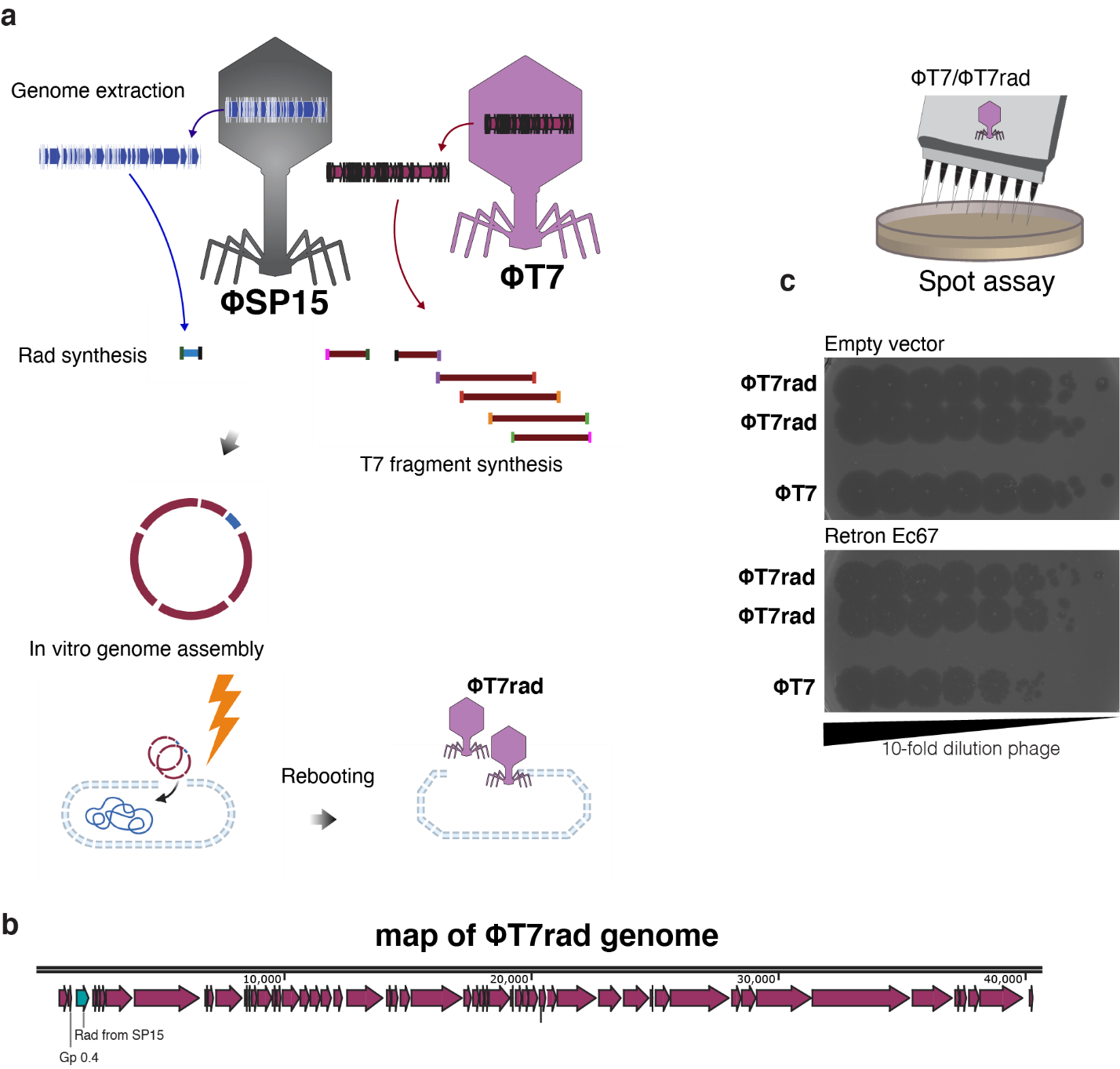

**Extended Figure 9.**

(a) Construction of *rad*-loaded T7 phage (T7rad) using synthetic approach. (b) Genome map of T7rad showing the *rad* gene from SP15 was introduced into the T7 genome downstream of *gp0.4*. (c) Spot assay of T7 and T7rad on bacteria carrying retron Ec67.

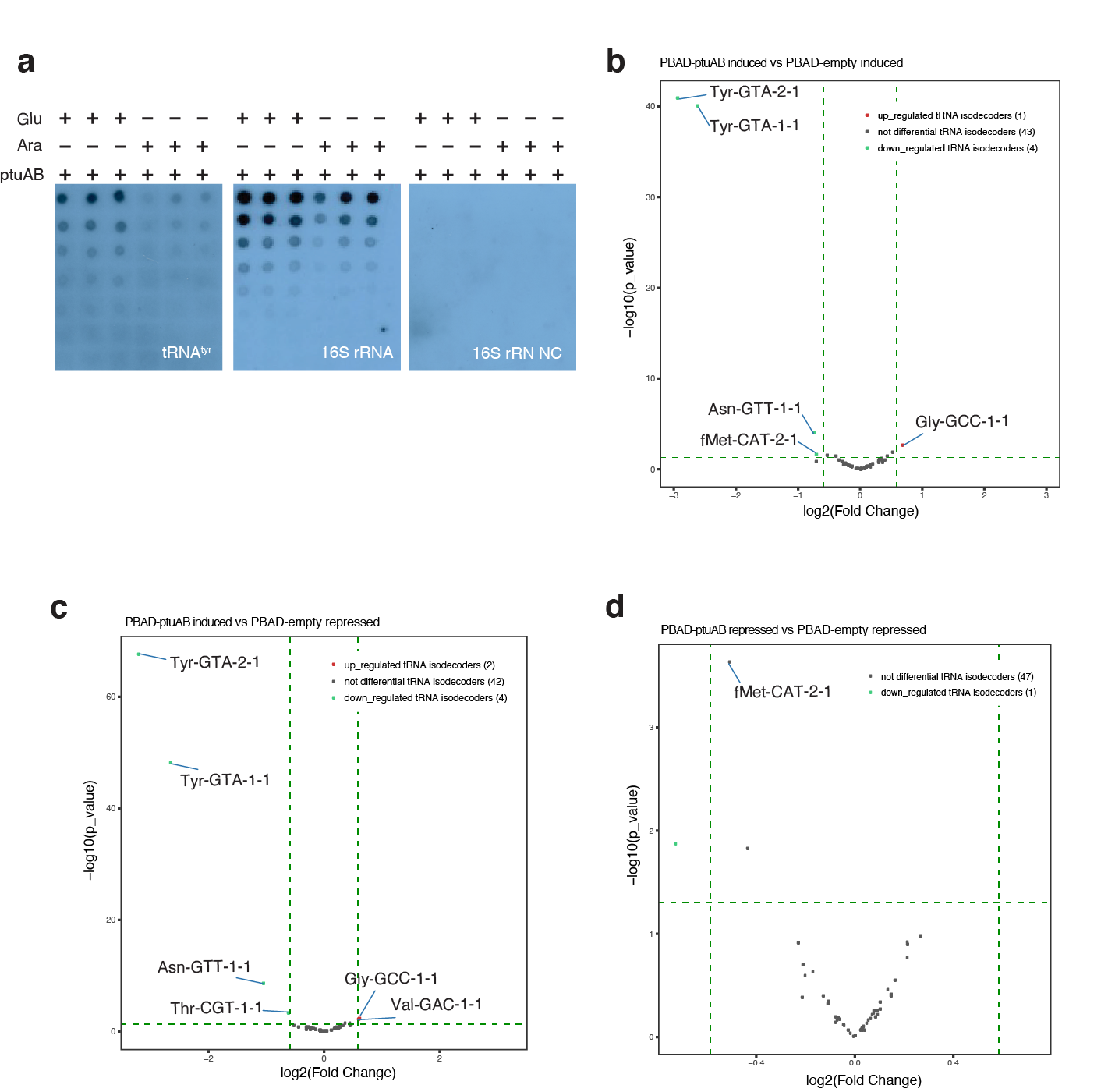

**Extended Figure 10.**

(a) RNA hybridization dot blot assay of bacteria that express PtuAB. From left to tight; tRNATyr, 16S rRNA, and 16S rRNA negative control. Sense oligo of 16S rRNA was used in negative control. (b) Expression level comparison between bacteria with induced PtuAB vs induced empty vector. (c) Expression level comparison between bacteria with induced PtuAB vs repressed empty vector. (d) Expression level comparison between bacteria with repressed PtuAB vs repressed empty vector. Plasmid expressing PtuAB or an empty vector was introduced into bacteria. Induction and repression of PtuAB-expression were performed with 0.2% of arabinose and glucose, respectively.

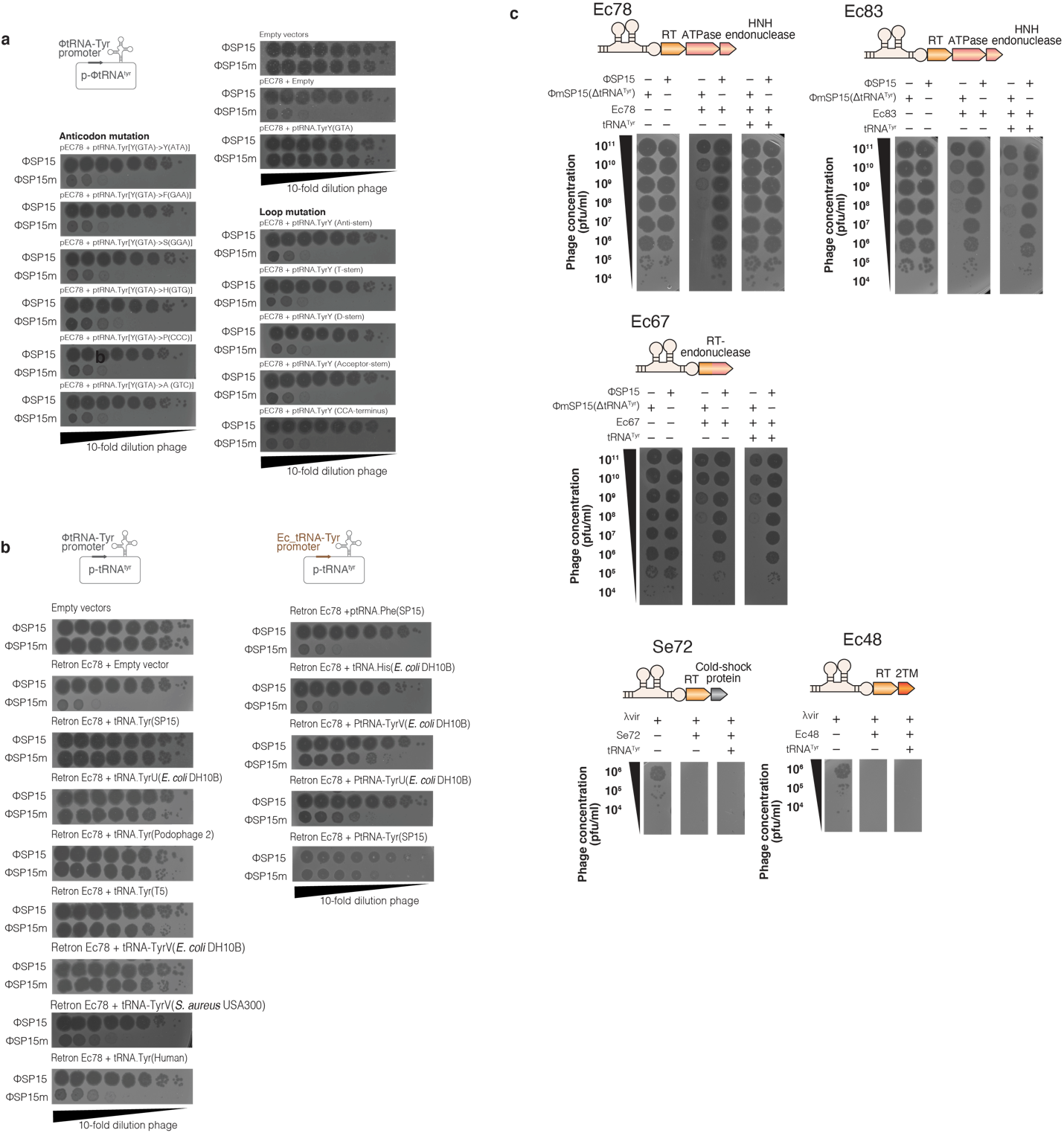

**Extended Figure 11.**

(a) Spot assay of phages, SP15 and its mutant SP15m on bacteria that carries retron Ec78 and various tRNA_Tyr_SP15 mutants. (b) Spot assay of phages on bacteria that carries retron Ec78 and various tRNAs from different organisms. (c) Spot assay of phages on bacteria carrying tRNA_Tyr_SP15 co-expressed with different type of retrons.

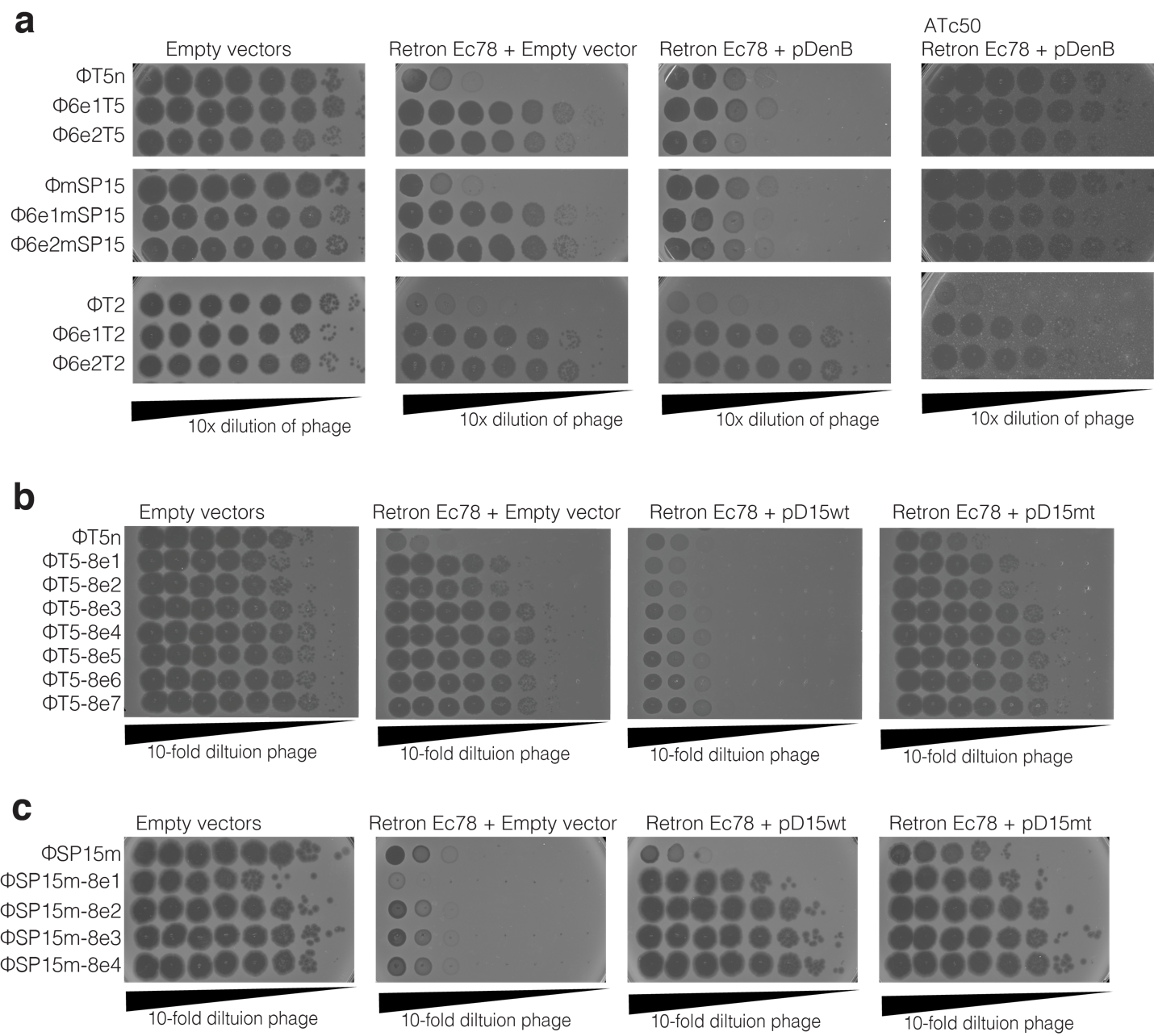

**Extended Figure 12.**

Spot assay of T5n phage and the escaper T5n mutant. (a) Spot assay of T5n, SP15m, and T2 and their respective escaper mutants on bacteria carrying retron Ec67 and DenB protein. SP15 and the escaper SP15 mutant (b) on bacteria carrying retron Ec78 and D15 protein. Complementation of D15 wildtype protein was performed in trans using plasmid pD15wt. Plasmids with mutant D15 (pD15m) was used for comparison.
